# FIND: Identifying Functionally and Structurally Important Features in Protein Sequences with Deep Neural Networks

**DOI:** 10.1101/592808

**Authors:** Ranjani Murali, James Hemp, Victoria Orphan, Yonatan Bisk

## Abstract

The ability to correctly predict the functional role of proteins from their amino acid sequences would significantly advance biological studies at the molecular level by improving our ability to understand the biochemical capability of biological organisms from their genomic sequence. Existing methods that are geared towards protein function prediction or annotation mostly use alignment-based approaches and probabilistic models such as Hidden-Markov Models. In this work we introduce a deep learning architecture (**F**unction **I**dentification with **N**eural **D**escriptions or **FIND**) which performs protein annotation from primary sequence. The accuracy of our methods matches state of the art techniques, such as protein classifiers based on Hidden Markov Models. Further, our approach allows for model introspection via a neural attention mechanism, which weights parts of the amino acid sequence proportionally to their relevance for functional assignment. In this way, the attention weights automatically uncover structurally and functionally relevant features of the classified protein and find novel functional motifs in previously uncharacterized proteins. While this model is applicable to any database of proteins, we chose to apply this model to superfamilies of homologous proteins, with the aim of extracting features inherent to divergent protein families within a larger superfamily. This provided insight into the functional diversification of an enzyme superfamily and its adaptation to different physiological contexts. We tested our approach on three families (nitrogenases, cytochrome *bd*-type oxygen reductases and heme-copper oxygen reductases) and present a detailed analysis of the sequence characteristics identified in previously characterized proteins in the heme-copper oxygen reductase (HCO) superfamily. These are correlated with their catalytic relevance and evolutionary history. FIND was then applied to discover features in previously uncharacterized members of the HCO superfamily, providing insight into their unique sequence features. This modeling approach demonstrates the power of neural networks to recognize patterns in large datasets and can be utilized to discover biochemically and structurally important features in proteins from their amino acid sequences.

**Author summary:** 

## Introduction

A central idea of molecular evolution is that homologous proteins in different biological organisms possess similar structural and functional properties. Homology of proteins is based on their amino acid sequences, which have some structural and functional properties encoded into them. The extent of sequence similarity of proteins can be used to classify proteins as belonging to different groups that each perform distinct functions or possess a specific set of properties. The proteins which fall within previously identified clusters are then annotated as having a specific function, which is usually experimentally determined for one or more proteins within a cluster. Computational tools such as sequence alignment based methods have been used to infer the biological roles of proteins, based on their similarity to experimentally characterized proteins [1]. These methods can be effective when comparing sequences with a high degree of similarity (60 percent or higher) but fail for more distantly related proteins [2]. Improvements were made to such methods by adding some position specific information to make local alignments [3], or by measuring sequence similarity against databases such as Pfam, with curated domain assignments [4]. Other methods include probabilistic models such as HMMer which calculate a sequence profile indicating the frequency of occurrence of a given amino acid in each position of a protein sequence [5], models incorporating information from phylogenetic trees such as SIFTER [6] or neural network based models where proteins are trained based on InterPro or GO classifications [2]. All of the above methods are more accurate for certain functions and biological systems, and they are evaluated regularly using the critical assessment of functional annotation (CAFA) challenge [7]. There are different challenges for each of the previous methods, some of which have been empirically demonstrated - BLAST alignments for example, cannot easily distinguish between parts of the sequence that align to conserved domains and some variable and less functionally relevant parts of sequences, and HMMs are more useful for annotation when built off a large data set for each protein domain. Neural networks offer some advantages over these methods.

Neural networks are computationally different from traditional methods, which are based on sequence alignments. Sequence alignments are computationally limited, and align sequences in a pair-wise manner and iterate over all possible pairs of sequences to compute the extent of similarity between proteins in a given database. In contrast, neural networks can process a larger set of sequences simultaneously to model the similarity in their amino acid sequences. With such advantages, neural networks have been shown to perform classification tasks with a high level of accuracy, even with smaller training sets [8] [9] [10] [11]. Another strength of neural networks is their ability to identify patterns or signals in large sets of data. Large protein superfamilies constitute such a large data set, from which it is conceivable that a neural network can identify “patterns” or functional features that correlate with functional characteristics, provided we curate a database with such specific and useful labels. An often cited disadvantage of neural networks is that they provide little information on the features that were used for classification. We have overcome this disadvantage by making our neural network model interpretable.

Our neural network model uses a neural attention mechanism to identify sequence features that are correlated with protein annotation. We use as input, a database of primary amino acid sequences that have been assigned “labels” that are defined based on phylogenetically identified clusters. The network is then trained on these protein sequences and labels to generate a classification model. This model is then used to provide annotation to a test set of sequences. The neural attention mechanism forces the model to use small sets of a protein’s sequence to classify it, thus identifying small functional or structural motifs that are inherent to a protein and correlated with the label it is assigned. While traditional methods require rigorous analysis of sequence alignments, extensive knowledge of prior biochemical literature and biochemical intuition, use of interpretable neural network models such as ours offers a useful time-saving tool. Direct extraction of sequence features unique to protein families will aid in mechanistic understanding of proteins unique to different organisms and pathways. Many attempts have been made to use certain characteristics such as % conservation, solvent accessibility or information on neighboring residues to correlate structural features with a protein’s function [12] [13] [14] but, our model is able to identify conserved sequence features directly from primary sequence and without being fed such curated information.

To evaluate our method, we built models to classify three different enzyme superfamilies – cytochrome *bd*-type oxygen reductases, nitrogenases and heme-copper oxygen reductases. We achieved a classification accuracy greater than 97 % for all of these families; comparable to a classification model we built based on Hidden-Markov Models. In addition, our model extracts structurally and functionally important sequence features. We compare the strengths and weaknesses of the above two approaches and discuss different parameters that can be optimized to enhance classification by the neural network. Further, we focused on and evaluated sequence features identified in the model as belonging to each protein family of the heme-copper oxygen reductase (HCO) superfamily. HCOs are a large superfamily of enzymes with several biochemically well-characterized family members. By using the case of experimentally characterized HCO enzymes, we validated our algorithm. Then, we utilized the model to discover several new interesting features in previously uncharacterized members of the HCO superfamily. The HCO superfamily classification model can be used to investigate new HCO sequences – to automatically identify the family they belong to, and extract novel features characteristic of that protein. Finally, our neural network approach is applicable to many protein databases, either small or large in dataset provided they are curated with appropriate labels to train on. When used with different protein superfamilies, we gain additional insight into their biochemical characteristics and into their evolutionary diversification. In order to make our tool accessible, we have made **FIND** available as a Jupyter notebook, allowing users to train any curated and labelled database of their choice for classification and feature extraction. Given the abundance of genomic and metagenomic sequence data that is currently available and the dearth of functional information, this tool will be helpful in generating additional insight into the possible role of divergent protein sequences as they are discovered.

## Heme-copper Oxygen Reductase Superfamily Background

The heme-copper oxygen reductase superfamily consists of families of respiratory enzymes – Oxygen reductases (*O*_2_-reductases) which perform oxygen reduction to water, or nitric oxide reductases (NOR) which perform nitric oxide reduction to nitrous oxide [15] [16]. It was hypothesized that one family (NOD) performs nitric oxide dismutation [17]. Briefly, these transmembrane enzymes receive electrons from a periplasmic or membrane bound electron donor (cytochrome c or quinol respectively) and use these electrons to perform oxygen or NO reduction. The core architecture of this enzyme typically constitutes a 12-transmembrane helix subunit where the active site is present (subunit I) and an electron-donor binding subunit where electrons are received from the physiological electron donor (subunit II). However, in one of the families (qNOR), these two subunits are fused into a single polypeptide. The central subunit I consisting of 12 transmembrane helices includes several features, conserved across the families that aid in catalyzing the reduction of *O*_2_ or NO. These features can be categorized in terms of protein components needed for (1)electron transfer to the active site, (2)proton transfer to the active site, (3)gas diffusion to the active site and (4) stabilization of redox-active components during catalysis. Co-factors which aid electron tranfer within HCOs include a high-spin heme that transfers electrons from subunit II to the active site and, the active site constituting a low-spin heme and a metal ion. Multiple sequence alignments (MSAs) and experimental studies have identified the amino acids which bind the above mentioned hemes and metals [16].

For proton transfer to the active site, there are conserved proton channels within some of the families, which (i) uptake protons from the cytoplasm for oxygen reduction to water and (ii) translocate protons from the cytoplasm to the periplasm (pumping protons) for generation of proton-motive force(pmf) [18]. This latter mechanism for generation of pmf is absent in cNOR and the proton channel leading from the periplasm replaces the cytoplasmic channel [19]. A gas diffusion channel for diffusion of *O*_2_ or NO to the active is present in all of the characterized HCO enzymes, with variations depending on whether the substrate is *O*_2_ or NO and the substrate affinity of a given enzyme [20] [21].

Finally, the two hemes involved in catalysis are stabilized by conserved arginines in the A and B-type *O*_2_ reductases, and by a Ca^2^+ in C-type *O*_2_ reductases and cNOR. The latter Ca^2^+ is stabilized by interactions with two glutamic acid residues. These glutamates are further stabilized by hydrogen bonding interactions with conserved residues, including a conserved arginine in both families [22] [19]. A fundamental difference in the active site of *O*_2_ reductases and NORs involves a unique co-factor – a conserved histidine and a tyrosine, found in all *O*_2_ reductases make a crosslink to each other and provide an electron to oxygen during catalysis [23]. The tyrosine from the cross-link is absent in the NORs and is replaced by a glutamate or asparagine in the experimentally characterized NORs [24] [25] [26]. All of the above mentioned features are detailed in **Figure 2** and **Table 1**.

**Table 1.** Conserved features in subunit I of heme-copper oxygen reductases, which are important for binding of heme or metal co-factors, transport of substrates to the active site, or catalysis of oxygen reduction to water

| Functional characteristic | Location in polypeptide sequence |
| --- | --- |
| Low spin heme binding site | Helix 2 and Helix 10 |
| High spin heme binding site | Helix 10 |
| $Cu_B$ or $Fe_B$ binding site | Helix 6 and Helix 7 |
| $O_2$ diffusion channel | Helix 2<br>and periplasmic loop between Helix 3 and 4 |
| D proton channel<br>(present only in the A-family) | Helix 2, Helix 3, Helix 4,<br>and cytoplasmic loop between Helix 2 and 3 |
| K proton channel | Helix 6 and Helix 8 |
| Stabilization of hemes (in C, cNOR) | Periplasmic loop between Helix 1 and Helix 2 |
| Stabilization of hemes (in A, B) | Periplasmic loop between Helix 11 and Helix 12 |

**Fig 2.**
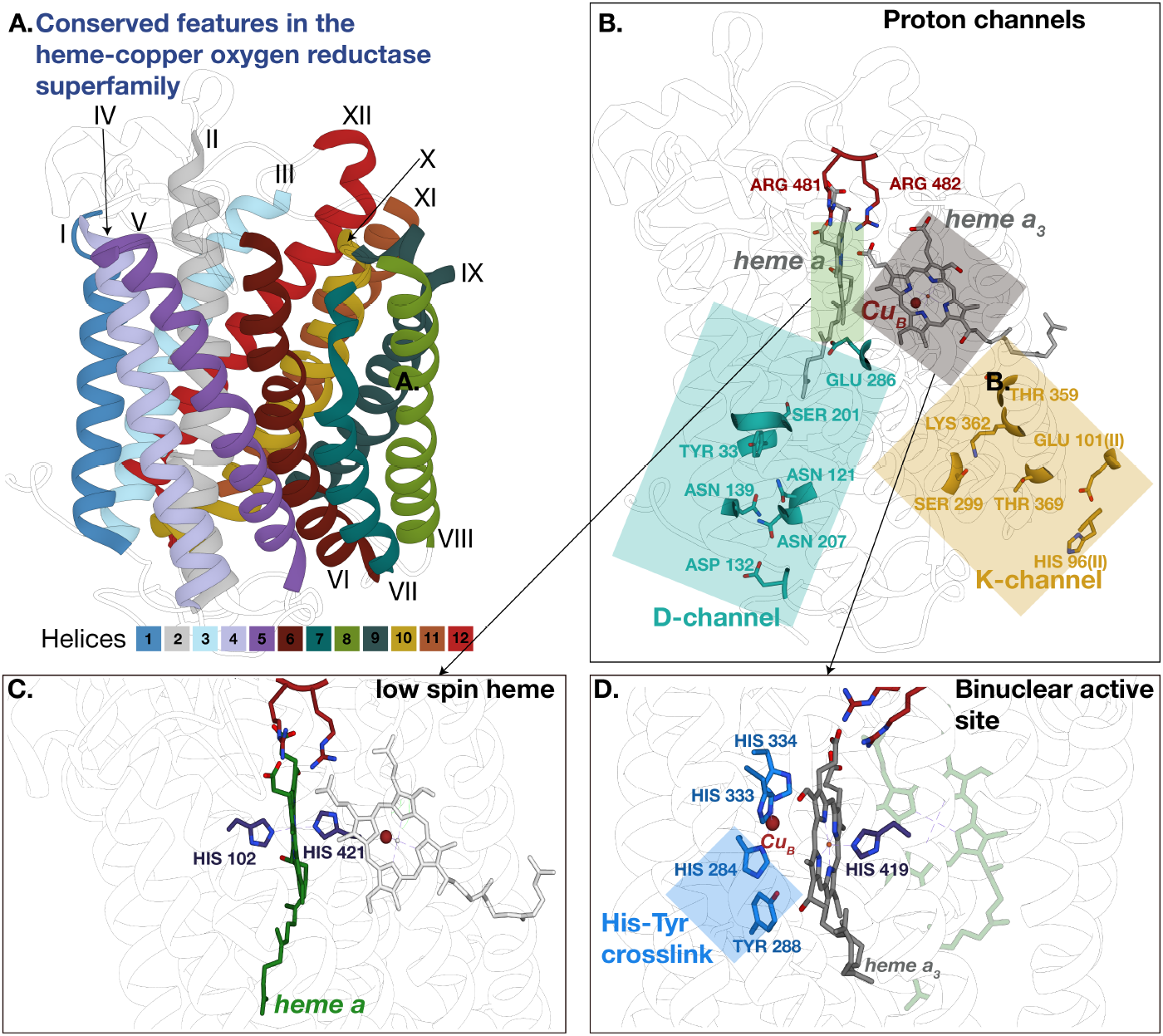
Conserved features in subunit I of the heme-copper oxygen reductases. These features, mentioned in **Table 1** were mapped onto the crystal structre of the A-family enzyme from *Rhodobacter sphaeroides* RCSB ID:2gsm.Panels **A** and **B** present identical views of the enzyme locating structural features within the context of their transmembrane helices. **A**. Structure of A-family enzyme color-coded according to the location of transmembrane helices in the enzyme. **B**. Location of the active-site heme and metal co-factors as well as conserved D- and K-proton channels within the HCO enzyme. **C**.Conserved histidines binding the low-spin heme in HCO. **D**.Conserved amino acids in the binuclear active site including the histidine-tyrosine crosslink that is unique to HCO.

To classify protein families within our model, only subunit I was used as was done for their original assignment using phylogenetics [15] **Figure 1**. 11 *O*_2_-reductase families and 7 NOR families were used to train the model for classification. NODs were left out of the model because of the smaller set of sequences available for this family. 8 of the *O*_2_-reductase families were previously described [15] while the I,J,K families have not been previously reported. The cNOR, qNOR and bNOR^1^, eNOR and sNOR [15] families have been previously described and so has the gNOR family [27]. The remaining family, nNOR is being reported for the first time but will be described in detail in a separate study^2^. A number of the conserved features mentioned above, as well as variations unique to families within the superfamily were discovered by the model.

**Fig 1.**
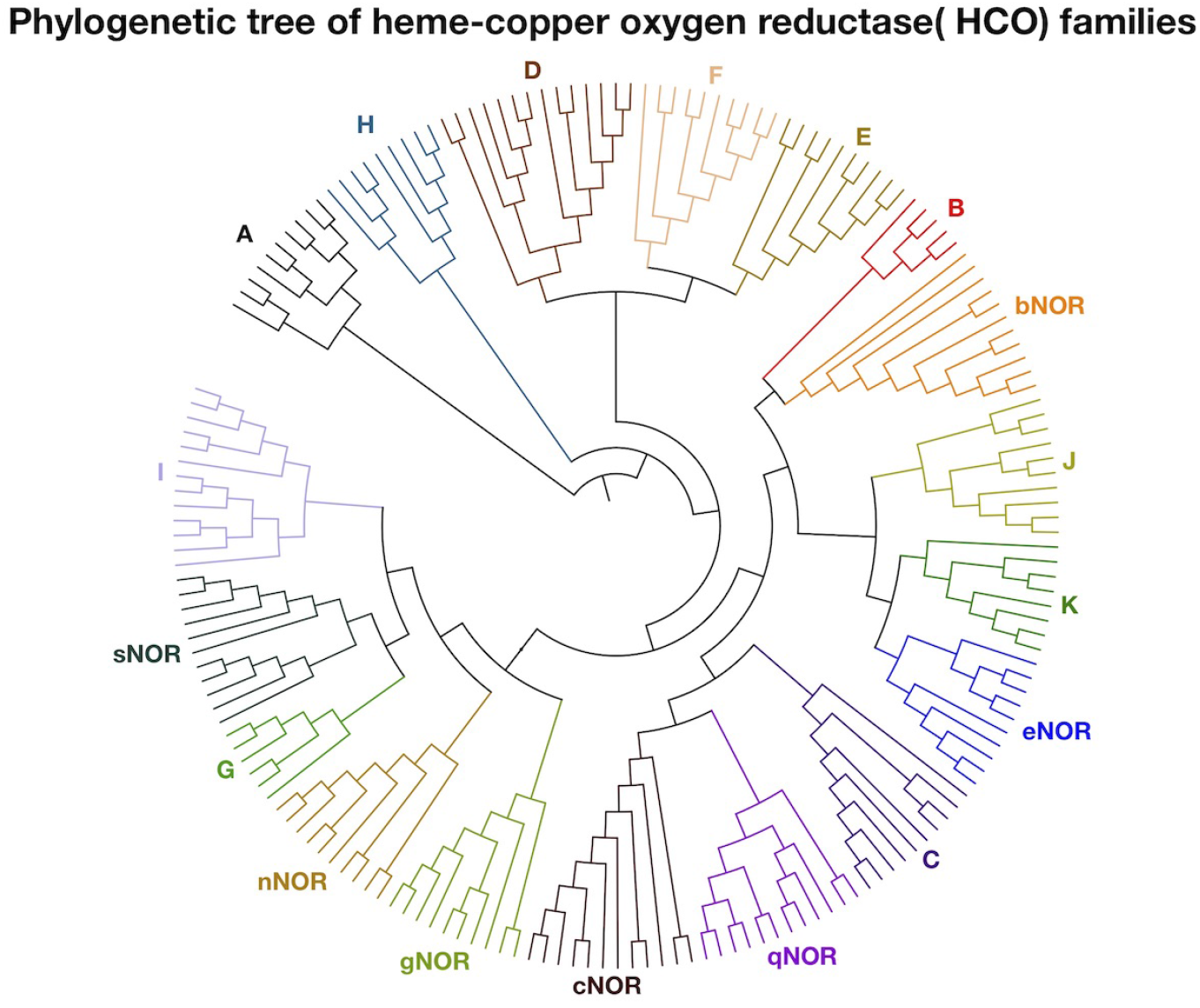
The heme-copper oxygen reductase superfamily is divided into 18 families based on monophyletic clades recovered through phylogenetic clustering of HCO amino acid sequences. The same clades were recovered using PhyML and MrBayes. The base sequence alignment used for the generation of this tree, as well as the phylogenetic trees are available on our Github database https://github.com/ybisk/FIND.

## Materials and Methods

### Data

Our model represents all proteins as a string of characters. Sequences from within the heme-copper oxygen reductase superfamily were used as input, with their functional annotation assigned by protein phylogenetics. The number of sequences within each family/class are reported in **Table 2**.

**Table 2.** Database of protein sequences used in this study from the following superfamilies - heme-copper oxygen reductase (HCO), nitrogenase, cytochrome *bd*. These sequences were split into a training set (**Tr**), validation set(**Val**) and test set(**Te**) for training the artificial neural network for protein classification, and for testing the accuracy of the model, respectively.

| heme-copper oxygen reductase |  |  |  | cyt *bd* |  | nitrogenases |  |
| --- | --- | --- | --- | --- | --- | --- | --- |
| Tr/Val/Te |  | Tr/Val/Te |  | Tr/Val/Te |  | Tr/Val/Te |  |
| A | 2569/384/729 | bNOR | 41/ 5/ 8 | A | 47/ 6/ 21 | Anf | 15/ / 5 |
| B | 535/ 87/160 | cNOR | 98/ 19/ 24 | B | 33/ 1/ 7 | nifDI | 164/ 28/ 61 |
| C | 819/126/229 | eNOR | 209/ 33/ 55 | C | 206/24/ 51 | nifDII | 107/ 15/ 21 |
| D | 66/ 9/ 15 | gNOR | 46/ 5/ 15 | E1 | 373/58/118 | nifD_ANME | 4/ 1/ 4 |
| E | 41/ 11/ 11 | nNOR | 33/ 4/ 7 | E2 | 123/17/ 33 | VnfD | 8/ / 3 |
| F | 26/ 4/ 8 | qNOR | 284/ 32/ 74 | E3 | 26/ 8/ 10 |  |  |
| G | 13/ 2/ 3 | sNOR | 140/ 23/ 40 | E4 | 41/ 3/ 9 |  |  |
| H | 13/ 1/ 8 |  |  |  |  |  |  |
| I | 17/ 1/ 3 |  |  |  |  |  |  |
| J | 11/ 1/ 3 |  |  |  |  |  |  |
| K | 5/ 1/ 1 |  |  |  |  |  |  |

### Computational Model (FIND)

In this section we introduce our model for **F**unction **I**dentification with Neural **D**escriptions or **FIND** for short. Our task is to predict the function of a protein based solely on its amino acid sequence. For this task, we use prior information in the form of a database of protein sequences which are correlated with a functional label. We then generate a computational model to identify parts of sequence features in the database sequences that correlate with its functional label. Then, the model is used to apply one of the previously curated functional labels to the query amino acid sequence by identifying the predictive features inherent in them. In computational terms, we need to find predictive regions of strings for a multi-class prediction which we train with gradient descent and a standard cross-entropy loss. Our task presents two unique challenges: long sequences and a comparatively small dataset.

The traditional approach to processing strings for categorization within Natural Language Processing is to use a Recurrent Neural Network (RNN) [28] sequence model to compress the input. Because our sequences range from 232 to 1095 characters, averaging over 500, RNNs, even when used with an long short-term memory (LSTM) cell [29], will suffer from vanishing gradients [30] and fail to learn. Secondly, as our goal is the construction of a model which provides interpretable insights into functionally predictive aspects of the input, we want to build an attention mechanism [31] which does not dramatically increase our parameter space, given the limited data we have access to. We are able to overcome both of these issues efficiently by using a weighted sum of local predictors, each resulting from a shallow stack of convolutions, and normalized confidence.

We present the basic structure of our model in **Figure 3**. A stack of convolutions [32] are applied until we reach a kernel of width 16 (*k* = 16). The final hidden vector is then passed through two feed-forward layers in order to produce both an attention weight and a set of predictions. The attention weights are normalized across the sequence and then multiplied by their corresponding prediction distributions (logits). This provides a single reweighted logit for the whole sequence scaled by each sub-predictor’s confidence. Finally, these values are summed and used for prediction at inference time or updated via a cross-entropy loss during training.

**Fig 3.**
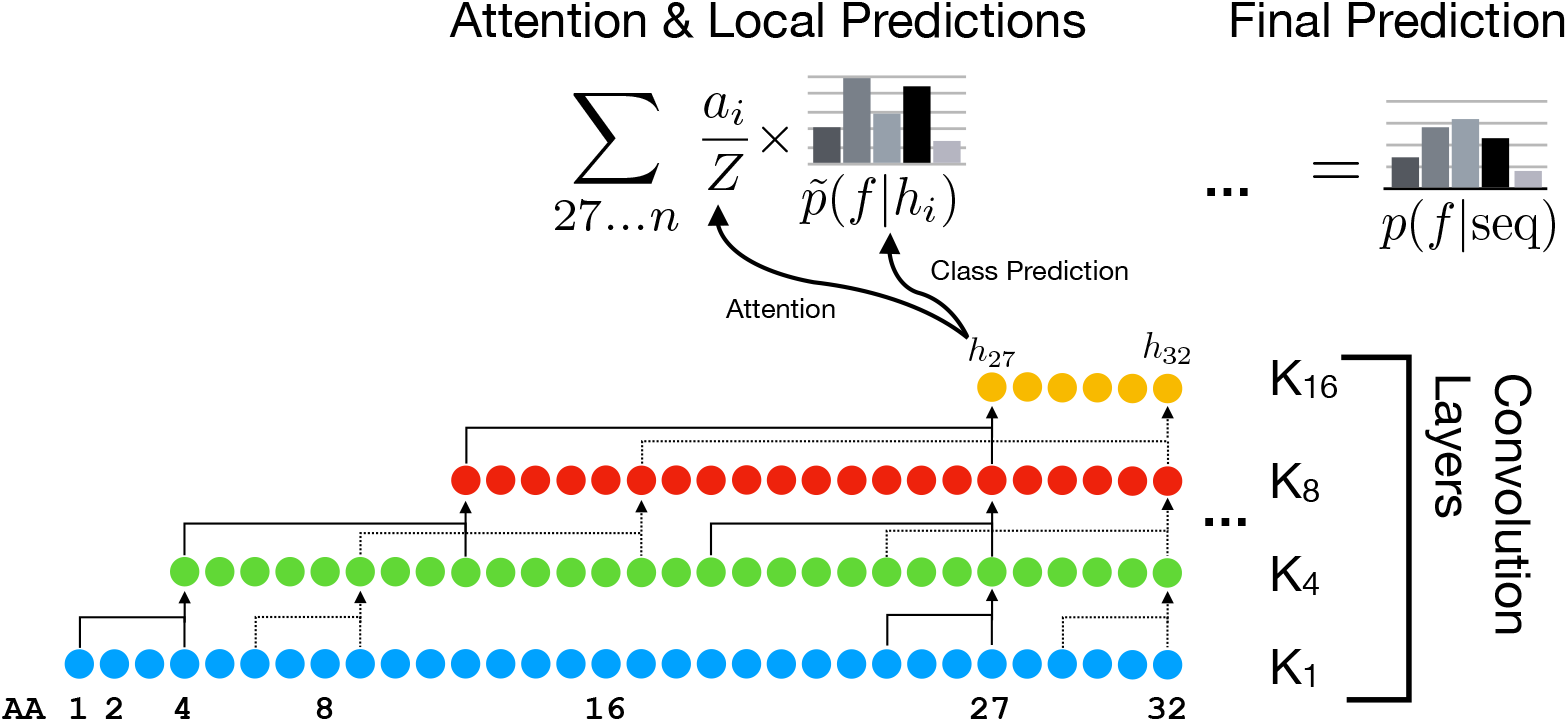
Our **FIND** model works by feeding Amino Acid sequences through increasingly wide convolutions (1, 4, 8, 16) yielding a final receptive field of 26. *h_i_* is multiplied by two fully connected layers to produce an attention weight and a prediction over the labels. The solid and dashed lines indicate example receptive fields for *h*_27_ and *h*_32_.

More formally, each unique amino acid type is assigned a random vector in 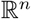, represented simply as an embedding matrix 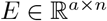. An input sequence *S* is therefore embedded as *e = S · E*, before passing *e* through 1-dimensional convolutions with kernel widths 2, 4, 8, and 16. This produces a final representation of the sequence 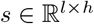 for a sequence of length *l*. Each element is then passed through a single linear layer to produce an attention (*α_i_ = s_i_ · A* where 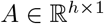) and prediction logits (*p_i_ = s_i_, × P* where 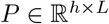 for *L* labels), *α* is then normalized via softmax to produce a distribution over all *l_i_* so the global sequence logits are simply computed as 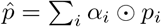, yielding a prediction vector 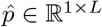, which is again normalized to form a distribution over the labels.

By normalizing attention across the entire sequence, the model is forced to choose a small set of sequences from which to make its prediction. This provides us with interpretable sequences to analyze. Finally, the convolution allows for efficient parallel analysis of every subsequence. All models were implemented in PyTorch,^3^ optimized with Adam [33], BatchNorm [34], and using a hidden dimension of *n* = 256 which was chosen based on the validation set performance. All code and data are available on GitHub for reproducibility and extension at https://github.com/ybisk/FIND. The code is also available in a Jupyter notebook for easy execution.

### Building a database for each enzyme superfamily

Most of the families for heme-copper oxygen reductase superfamily were previously defined in [15]. The previously undefined families of NOR were designated in the same way, creating phylogenetic trees and separating out enzyme sequences belonging to monophyletic clades within the tree. Once the families were defined,^4^ a query sequence for each of the families was used to search the NCBI non-redundant protein sequence database using Biopython’s Entrez module and the BLAST tool to identify hits. Accession numbers for the hits were retrieved using the SeqIO module and then protein sequences were retrieved, again using Entrez to connect to the NCBI database. Bit Score and e-value cut-offs could not be identified to entirely separate the sequences according to families. Therefore, the sequences were then manually curated and labelled as belonging to their respective families. When enough sequences were not identified in the NCBI database, sequences were retrieved from metagenomes on JGI’s IMG server. All accession numbers are provided in Supplementary information. A similar method was used for building a database for cytochrome *bd*-type oxygen reductases and nitrogenases. However, no metagenome sequences were included for nitrogenases. The nitrogenases database also included unpublished sequences from a private database.^5^

### Analysis of “predictor” sequence features learned by the model

The model was forced to choose small sections of a query sequence to classify a given protein as belonging to one of the different HCO families. The extracted sequence feature was then retrieved and represented using a reference crystal structure or model in the following way. An (MSA) was generated using the query sequence that the model has just classified, and a set of previously curated set of HCO sequences belonging to the output class. The extracted feature was searched against the MSA to locate the corresponding location in the sequence of the reference structure. The “predictor” sequences were then mapped using different colors onto the reference structure using Chimera. [35]

### Generation of structural models

Crystal structures for previously characterized enzymes within the HCO superfamily were used to display the sequence features predicted by the learning algorithm. When a crystal structure was not available, models of the HCO family were generated using the i-TASSER structural modelling server. [36] Reference sequences used for generating a model and the quality of the model itself are found in Table S2. The models themselves are available on Github https://github.com/ybisk/FIND.

### A Hidden Markov Model classifier for heme-copper oxygen reductases

Using the same test set as that defined for learning the classification model, we created MSAs for each of the families identified within the HCO superfamily. These MSAs were used to create a HMM profile for each protein family, using the HMMbuild function in the HMMer module [5]. A file with a set of query sequences was then searched against each of the HCO-family HMM profiles, using the hmmscan function. The family against which the query sequence achieved the highest domain-based Bit Score was taken to be its label. Classification accuracy was then determined using the standard parameters of precision, recall and the harmonic mean, F1.

## Results and Discussion

Our model was evaluated for its ability to classify a protein belonging to one of three test superfamilies - cytochrome *bd*-type oxygen reductases, nitrogenases and heme-copper oxygen reductases. Its accuracy at performing the above task was compared to that of a Hidden-Markov based classification tool, using standard evaluation paramaters. Thereafter, the sequence features used by a model for the HCO family members were validated using prior biochemical literature on the subject. The validated model was used to classify and characterize enzymes from the families that have not been experimentally tested.

### Comparison of classification accuracy – CNNs and Hidden Markov Models

In all three test cases, the CNNs performed classification with an accuracy greater than 96%.^7^(**Table 3**) This was comparable to the classification accuracy achieved by an HMM-based classifier that we designed. Impressively, the CNN achieves a high level of classification accuracy for the relatively small data set of nitrogenases, though the HMM classifier achieves perfect accuracy in that case. It is conceivable that HMM-based classifiers are more accurate for smaller, more homogenous data sets as there is less variation across a domain in such datasets. This might present a case of over-fitting the data, which is less likely in the case of CNNs which model all of the HCO sequences at the same time. In the cytochrome *bd*-type oxygen reductases, both the HMM and CNN perform classification with greater than 99% accuracy. Interestingly, HMM and CNNs are more accurate for different families within the large superfamilies. In HCO classification for example, the CNN predicts the H and nNOR families more accurately than the HMM, while the HMM is more precise when predicting several families, especially F and G. There are two fundamental difference between these approaches – the CNN uses smaller sections of sequences to make a prediction of function while the HMM uses whole domain similarity, and the CNN models all sequences simultaneously while the HMM uses pairwise comparisons. It is possible that our CNN model performs well where unique features, comparable to the length of our receptive field are easy to identify, while the HMMs may identify a more dispersed combination of sequence features. In the HCO superfamily, it is apparent that unique features are formed through evolutionary diversification and are conserved. Correspondingly, our model is able to easily classify members within the HCO family.

**Table 3.** Protein classification accuracy achieved using a convolution neural network in comparison to a Hidden Markov Model-based classifier. Classification accuracy is evaluated using – Precision (**Pr**), Recall (**Re**) and Harmonic mean (**F1**).^6^

| (a) HCOs |  |  |  |  |  |  |
| --- | --- | --- | --- | --- | --- | --- |
| HCO | CNN |  |  | HMM |  |  |
|  | Pr | Re | F1 | Pr | Re | F1 |
| A | 1.00 | 1.00 | 1.00 | 1.00 | 1.00 | 1.00 |
| B | 0.98 | 1.00 | 0.99 | 0.99 | 1.00 | 0.99 |
| C | 1.00 | 1.00 | 1.00 | 1.00 | 1.00 | 1.00 |
| D | 0.95 | 1.00 | 0.97 | 1.00 | 1.00 | 1.00 |
| E | 0.83 | 1.00 | 0.91 | 1.00 | 1.00 | 1.00 |
| F<sup>†</sup> | 0.80 | 1.00 | 0.89 | 1.00 | 1.00 | 1.00 |
| G<sup>†</sup> | 0.67 | 1.00 | 0.80 | 1.00 | 1.00 | 1.00 |
| H<sup>†</sup> | 1.00 | 1.00 | 1.00 | 0.89 | 1.00 | 0.94 |
| I<sup>†</sup> | 1.00 | 0.71 | 0.83 | 0.75 | 1.00 | 0.86 |
| J<sup>†</sup> | 1.00 | 1.00 | 1.00 | 1.00 | 1.00 | 1.00 |
| K<sup>†</sup> | 0.00 | 0.00 | 0.00 | 1.00 | 1.00 | 1.00 |
| bNOR | 1.00 | 1.00 | 1.00 | 1.00 | 1.00 | 1.00 |
| cNOR | 0.85 | 1.00 | 0.92 | 1.00 | 1.00 | 1.00 |
| eNOR | 0.96 | 1.00 | 0.98 | 1.00 | 1.00 | 1.00 |
| gNOR | 1.00 | 0.88 | 0.93 | 1.00 | 1.00 | 1.00 |
| nNOR<sup>†</sup> | 1.00 | 0.56 | 0.71 | 1.00 | 0.33 | 0.50 |
| qNOR | 0.99 | 0.95 | 0.97 | 1.00 | 1.00 | 1.00 |
| sNOR | 0.92 | 1.00 | 0.96 | 1.00 | 1.00 | 1.00 |
| Acc | 99.14% |  |  | 99.96% |  |  |

<sup>†</sup>Families with <10 sequences.

| (b) Nitrogenases |  |  |  |  |  |  |
| --- | --- | --- | --- | --- | --- | --- |
| Family | CNN |  |  | HMM |  |  |
|  | Pr | Re | F1 | Pr | Re | F1 |
| nifDI | 1.00 | 1.00 | 1.00 | 1.0 | 1.0 | 1.0 |
| nifDII | 0.96 | 1.00 | 0.98 | 1.0 | 1.0 | 1.0 |
| Anf | 1.00 | 1.00 | 1.00 | 1.0 | 1.0 | 1.0 |
| nifD_ANME | 1.00 | 0.25 | 0.40 | 1.0 | 1.0 | 1.0 |
| VnfD | 0.60 | 1.00 | 0.75 | 1.0 | 1.0 | 1.0 |
| Acc | 96.8% |  |  | 100% |  |  |

(c) Cytochrome *bd*-type oxygen reductases
| Family | CNN |  |  | HMM |  |  |
| --- | --- | --- | --- | --- | --- | --- |
|  | Pr | Re | F1 | Pr | Re | F1 |
| A | 1.00 | 1.00 | 1.00 | 1.0 | 1.0 | 1.0 |
| B | 1.00 | 1.00 | 1.00 | 1.0 | 1.0 | 1.0 |
| C | 1.00 | 1.00 | 1.00 | 1.0 | 1.0 | 1.0 |
| E1 | 0.99 | 1.00 | 1.00 | 1.0 | 1.0 | 1.0 |
| E2 | 1.00 | 1.00 | 1.00 | 1.0 | 1.0 | 1.0 |
| E3 | 1.00 | 0.90 | 0.95 | 1.0 | 1.0 | 1.0 |
| E4 | 1.00 | 1.00 | 1.00 | 1.0 | 1.0 | 1.0 |
| Acc | 99.6% |  |  | 100% |  |  |

<sup>†</sup>Families with <10 sequences.

The performance of the model on the HCO superfamily, particularly with families with small data sets is impressive. Previously, an HMM-based classifier was created for the HCO superfamily but, it does not incorporate as much phylogenetic diversity of this superfamily as we present here [37]. Therefore, we built an updated model for this task, which performs the task with >99 % accuracy. It is interesting to note that classification by both CNN and HMM is least accurate in the case of the I family of *O*_2_-reductases and nNOR, which is consistent with these being a more complicated task for both algorithms that we used. Further, it appears that the false positives for the I-family is often the G-family, which is corroborated by their close phylogenetic relationship (**Figure 1**). This is also true for nNOR and sNOR, which are closely related. Sequence analysis of subunits II from these enzymes also suggest that they have recently diverged, with respect to their evolutionary history. It is intriguing that even when we force the model to make predictions using smaller regions of the amino acid sequence, it is sensitive to close phylogenetic relationships. Our feature extraction method allows us to get a tangible understanding to the physical nature of these close evolutionary relationships, i.e what features are common to close evolutionary partners and which features vary between distant ones.

### Predictor sequences obtained using attention mechanism

One of the primary goals in this work is to identify inherent patterns within protein sequences that may not be easy to identify without prior knowledge and biochemical intuition. In accordance with that goal, we forced our classification algorithm to identify parts of the protein sequence that are most conserved within a given family of enzymes. The algorithm identifies these sections of protein sequences, or sequence features, by preferentially weighting those that are correlated with the correct functional or structural label. We validate these “predictor” sequence features against prior knowledge of the known HCO families, and then characterize the novel sequence features within the new families.

#### Validation of the model using extensively characterized HCO families

The classification model correctly identifies several features from the known HCO families - A, B, C, cNOR and qNOR (**Figure 4, Table S1**). From the A-family, the most often extracted feature is in the active site (present in Helix 6), including either the conserved glutamate or tyrosine and serine, which are the amino acids from which protons are loaded onto the oxygen molecule as it is reduced at the active site (**Figure 4**). One of these two variants is always present in the A-family and only in this family. The C-family “predictor” sequence most often identified is the tyrosine belonging to the histidine-tyrosine crosslinked cofactor, which is located on a different helix (Helix 7), than in the rest of the *O*_2_-reductases where it is found in Helix 6(**Figure 4**). In the B-family, the model derives motifs corresponding to its active site and a tyrosine that is conserved in the proton channel (**Figure 4**). In the cNOR family, one often identified feature is around Helix 10, including a conserved threonine residue suggested to line a proton channel leading from the periplasm to the active site [19]. (**Figure 4C.**). In both C and cNOR, a conserved arginine which is involved in stabilizing the active site heme is present. cNOR has a glutamate in place of the tyrosine from the cross-linked cofactor present in the *O_2_*-reductases. All the sequence features derived were consistent with previous experimental results. In fact, the identified locations agree well with those listed in **Table 1**, indicating that the model is extracting parts of a given enzyme’s sequence that correlate with meaningful aspects of its function.

**Fig 4.**
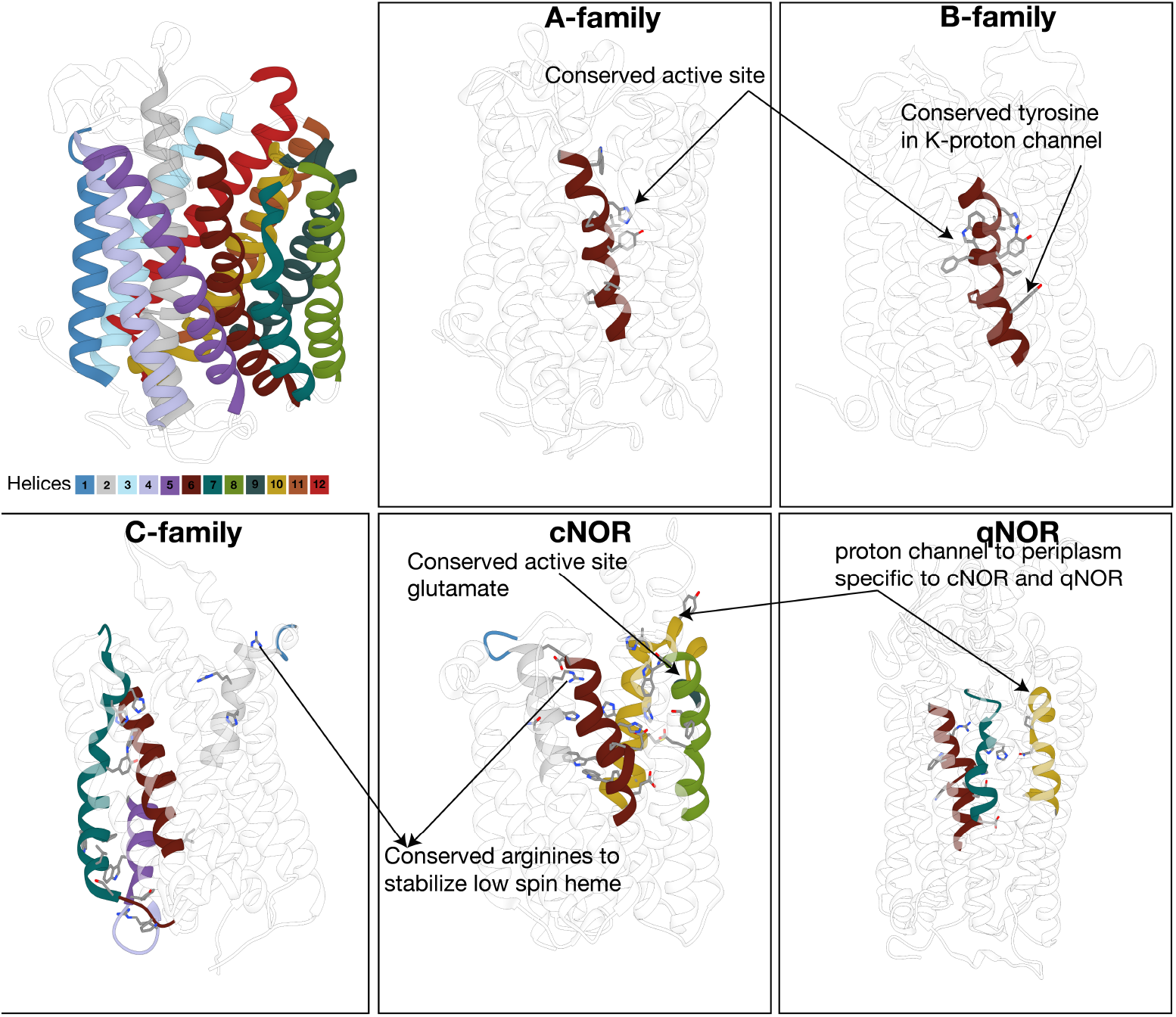
Sequence features revealed by classification algorithm. Sequence features were mapped onto crystal structures of HCO enzymes that are well characterized. The PDB codes for A, B, C, cNOR, qNOR family crystal structures used are 2gsm, 1xme, 3mk7, 3wfb, 3ayf respectively

#### Identification of sequence characteristics unique to previously uncharacterized HCO families

After validating the model, we used the model to extract sequence features from the remaining families which have not been well characterized. Characteristics unique to each family were uncovered, indicative of their adaptation to a unique environment and role. Among the more striking sequence traits was recognized in the J family, which has two tyrosines replacing histidines which are typically ligands to the active site heme and high-spin heme. (**Figure 5**) This is likely to have an effect on the midpoint potential of the heme; the anionic nature of tyrosine as a ligand is likely to lower the midpoint potential of the heme. [38] Also interesting are residues corresponding to a proton channel in the nNOR family in a structurally analogous location to that of the B family. This would be physiologically significant, indicating that this enzyme could generate proton motive force. A similar channel leading from the cytoplasm to the active site has been shown in the electrogenic bNOR [26]. nNOR also appears to have a histidine to methionine substitution for a ligand to the low-spin heme in the electron transfer pathway. A histidine to glutamate substitution as a heme ligand is uncovered in gNOR, which would have a significant effect on the midpoint potential and spin state of the heme. Finally, in K and eNOR, conserved residues around the *O*_2_/NO diffusion channel are found. The full set of features is listed in **Table S1** with corresponding amino acid numbering for previously characterized HCOs.

**Fig 5.**
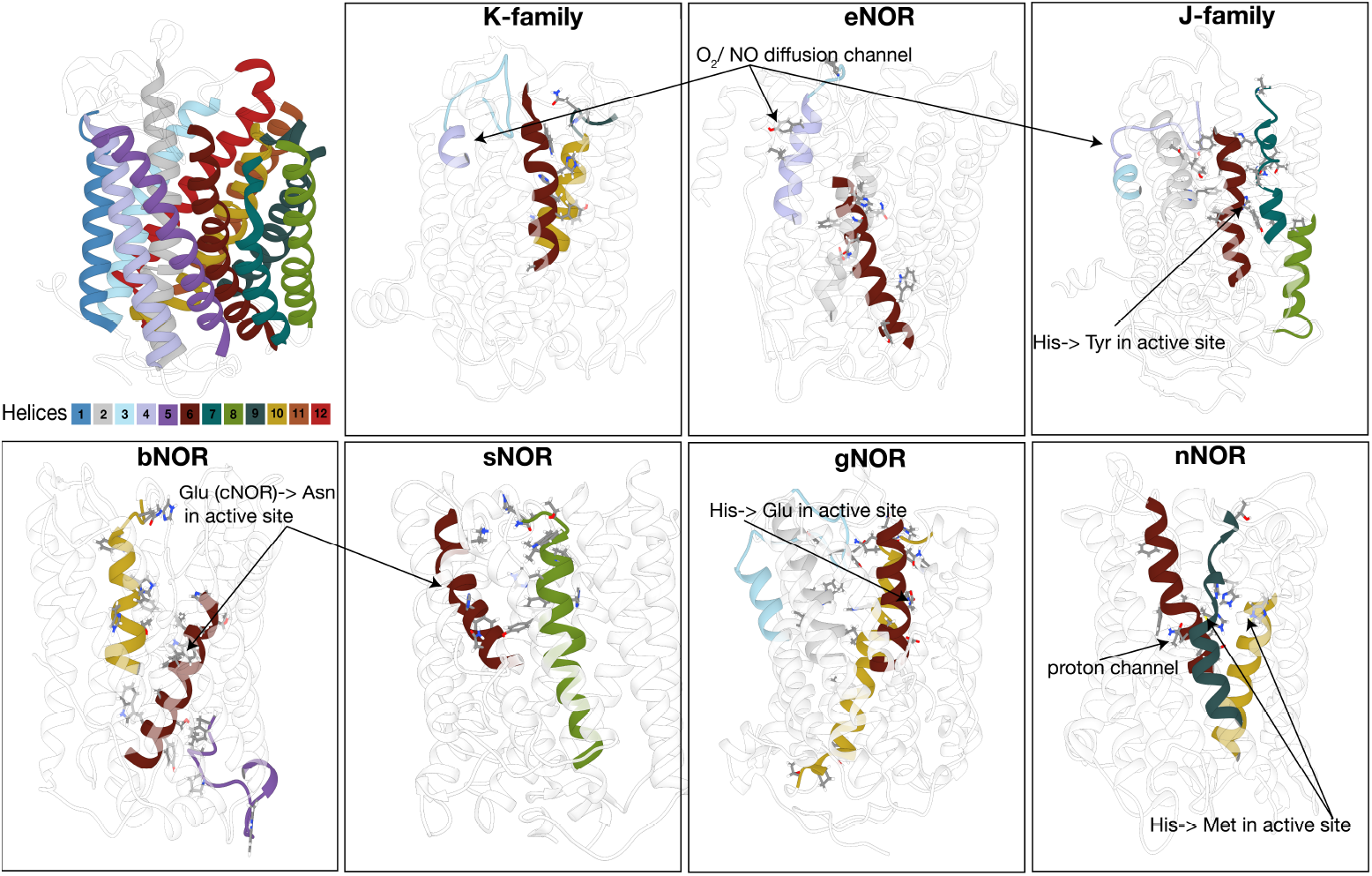
Sequence features revealed by classification algorithm. Sequence features were mapped onto homology models of HCO enzymes that are uncharacterized. The structural model and quality parameters for each model used for each family is identified in **Table S2**

By using these extracted features, it has thus been possible to gain insight into some functional and structural characteristics of biochemically uncharacterized proteins. It is interesting that most of the features that have been identified are part of transmembrane helices. This is consistent with a majority of the catalytically important residues in HCO enzymes being present in the membrane.

A brief analysis of the sequences extracted by the CNN from nitrogenases’ classification indicates that active site features are used as the markers for classification. This suggests that this method is as successful for smaller and cytoplasmic proteins. Extraction of features from cytochrome *bd* identify locations near the electron donor binding site, and certain periplasmic loops. A full description of features identified for nitrogenases and cyt bd is beyond the scope of this work and will be presented elsewhere.

### A simplified model of functional diversification within the HCO family members

#### Use of different model parameters to gain insight into different protein features

The neural network model is built on several parameters - the size of the embedding matrix, the number of convolutional layers and kernel size - that can be modified to improve classification accuracy as well as extract different features from protein sequences. The embedding matrix is used to encode each amino acid in a protein sequence with a unique vector that, in a sense, quantifies the characteristics associated with that amino acid. Modification of the the size of the embedding matrix used to code for an input sequence can modify the complexity available to the model. Intuitively, this would vary the model’s ability to perceive relationships between different amino acids, for e.g., between polar residues or between acidic amino acid residues. Further, the number of convolutional layers and kernel sizes varies the length of the receptive field, or essentially the maximum length of sequence the model is capable of attending to when using the attention mechanism. The attention mechanism is used to extract sequence features, and the model seems capable of using as small a receptive field as 15 amino acids to accurately predict classes of protein families. Use of dilation in the neural networks can increase this receptive field further and the most striking contrast in the features learned by the model is perceived when combining dilation with the number of layers used to encode complexity. Significantly, using weighting parameters to increase the importance of the rarer classes to the learning of the model, improves classification accuracy. A careful examination of model parameters used to build a neural network indicates that there is significance to the modified use of these parameters. (Data present in Github https://github.com/ybisk/FIND) However, the consistency in the features learned for the different classes using a weighted, convolutional network with 256 parameters in the embedding matrix is the highest and we used these parameters to generate the final model for the HCO classification.

Using the classification model learned by the neural network, we extracted the top “predictors” or weights assigned to various parts of the sequence within HCO family members. Simply, this provides us with sequence patterns or motifs correlated with a given label. These patterns were plotted using a heat map to demonstrate the location of identified sequence features within subunit I of the enzyme. We find that several of these patterns correlate with the evolutionary history of the HCO family, as shown in **Figure 6**. Comparison of the distribution of identified features within different families demonstrates the differential functional adaptation of some of the families. For instance, the regions identified as predictive in cNOR and qNOR center around helices 8, 9 and 10 where residues lining a proton channel for protons to enter the active site from the periplasm are found. Conversely, A, B and C families of *O*_2_ reductases have a proton channel leading from the cytoplasm to the active site. Sequence motifs found in the C-family oxygen reductase are centered around the region between helices 1 and 2, where a periplasmic loop exists that stabilizes heme *b_3_* in the C-family [22]. Conserved features are identified in cNOR from the same region. The above identified correlations are significant because they correspond to the phylogenetic relationship between the C-family *O*_2_ reductase, and cNOR and qNOR. The close evolutionary relationship between the C-family, cNOR and qNOR has been observed before [39] [25] and the common structural features associated with these families appear to corroborate that. A similar relationship is observed between K and eNOR which cluster closely together in our HCO phylogenetic tree. Conserved residues such as threonine and tyrosine (in the K-family), and a tyrosine in eNOR in the loop between Helices 3 and 4, likely affect the *O*_2_ and NO diffusion pathway, which have been explored in the A, B and C-family [21]. These features are suggestive of adaptations specific to substrate concentrations [40]. Within the context of a multi-class classification model, we intuit that structural differences between the families are weighted and extracted. This implies that the model does not discover every feature conserved in a family but uncovers those characteristics that are unique to each clade. A complete picture of each enzyme family’s catalytic and structural characteristics may not be obtained from the model but, it has the advantage of providing insight into the bigger picture of functional diversification within the enzyme superfamily. This in turn is correlated with the nature of catalytic machinery and structural features within a given family. In identifying features that are significant enough to be conserved within a clade and yet, adaptable within the larger superfamily, we discern what some of the constraints for evolution of these proteins are and how they diverged through time.

**Fig 6.**
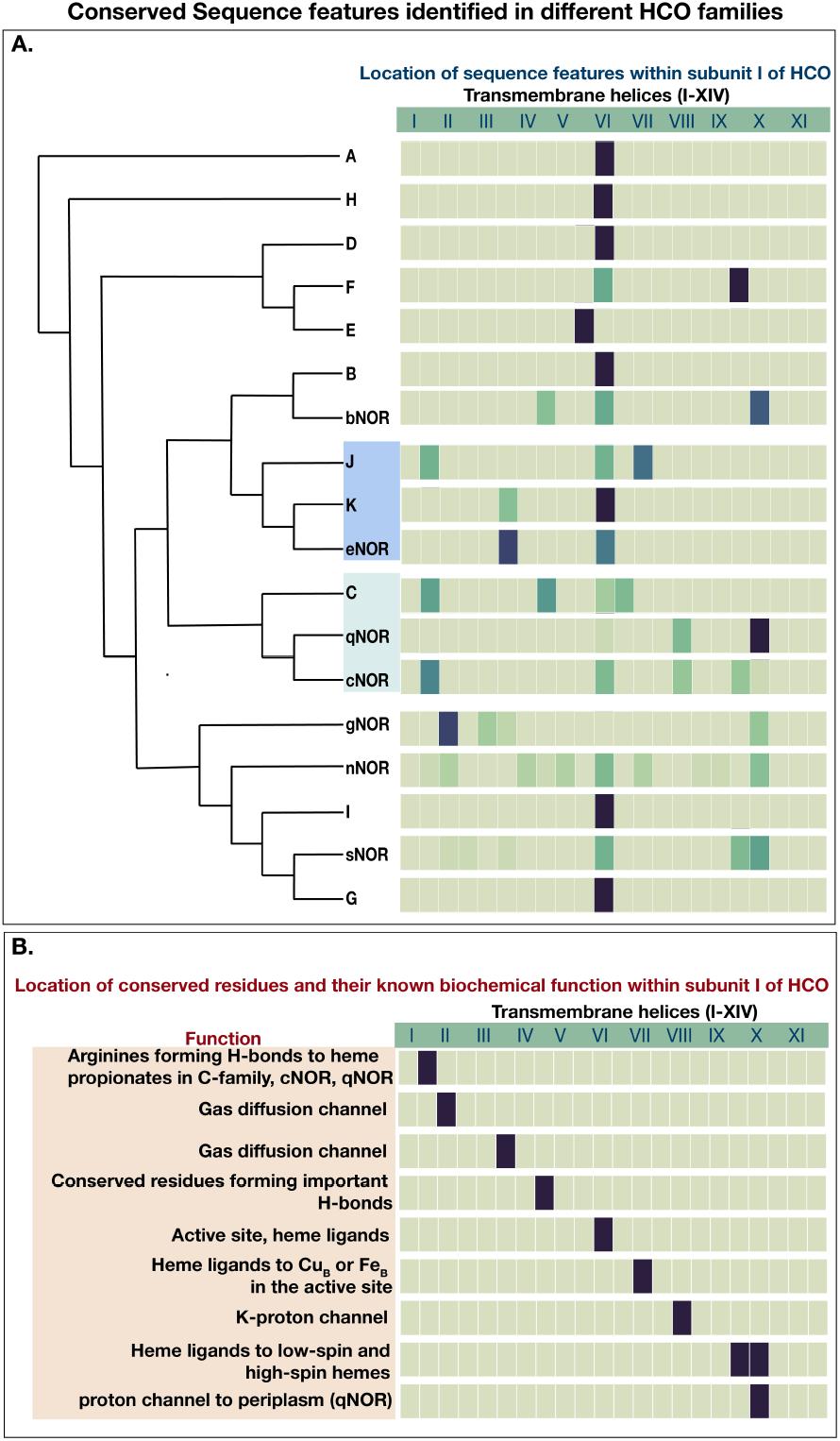
Correlation of evolutionary history with sequence features identified in different HCO families. A heatmap of extracted features as they are located within the conserved subunit I, is correlated with the phylogenetic relationships identified in **Figure 1**. Features from five different training runs were combined and mapped onto subunit **I**.

## Conclusion

Our **FIND** classification model has accurately classified members of three different enzyme superfamilies - heme-copper oxygen reductases, nitrogenases and cytochrome *bd*-type oxygen reductases. It performs this task with accuracy comparable to a well-accepted classification technique based on Hidden Markov Models. In addition, our model is able to automatically extract interesting, unique sequence features correlated with the labels we imposed. This facet of our model allows us to infer biochemically and structurally distinct features in protein sequences without extensive review of sequence alignments, which are typical of traditional methods. In this way, it is uniquely useful for investigating large sets of uncharacterized proteins. Further, our model provides insight into the evolutionary relationships of proteins within an enzyme superfamily, by identifying some traits that are specific to each protein. The novel uncharacterized members of heme-copper oxygen reductases that we have described and the unique features we have identified in their protein sequences would be an interesting set of targets for biochemical characterization. In our work, we have seen the promise of deep learning methods; to classify proteins and to identify innate structural and functional features associated with function, directly from primary sequence. In the future, these methods can be used to tackle the heterogeneity in biological sequence data and understand aspects of protein diversity and functional adaptation.

## Acknowledgments

This work was funding by DARPA’s CwC program through ARO (W911NF-15-1-0543), the Moore Foundation(Award to VJO: GBMF3780), the Department of Energy (Award to VJO: DE-SC0016469) and the Center for Dark Biosphere Investigations(NSF Award to VJO: OCE-0939564). We are grateful to Daan Speth for reading the manuscript at multiple stages of completion and for his useful comments throughout the process. We would also like to thank Dr. Robert Gennis for reading the manuscript and offering helpful comments, Grayson Chadwick for useful discussion on bioinformatics strategies and Haley Sapers for her instructive critique of the figures.

---

**Table S1.**
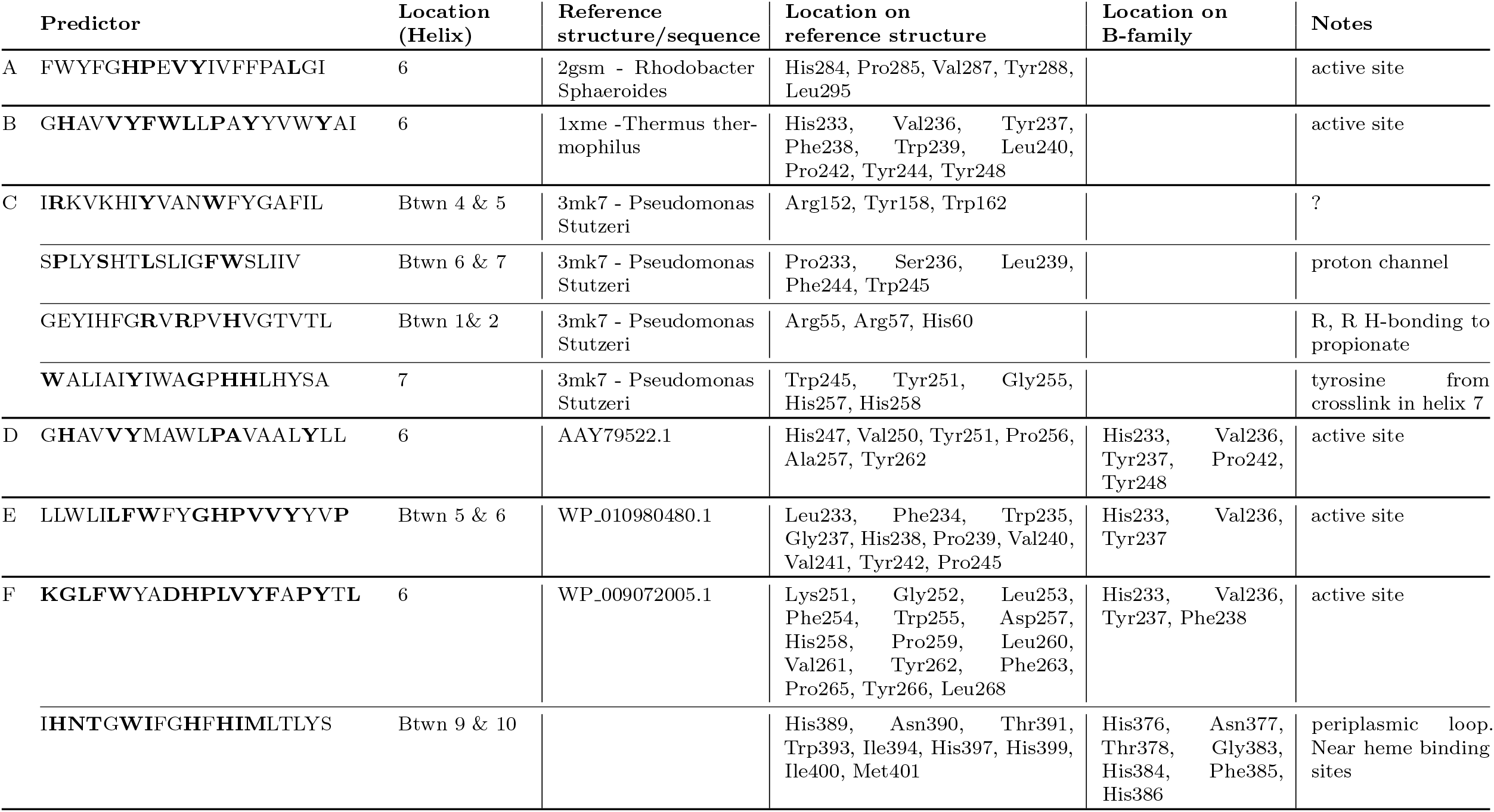
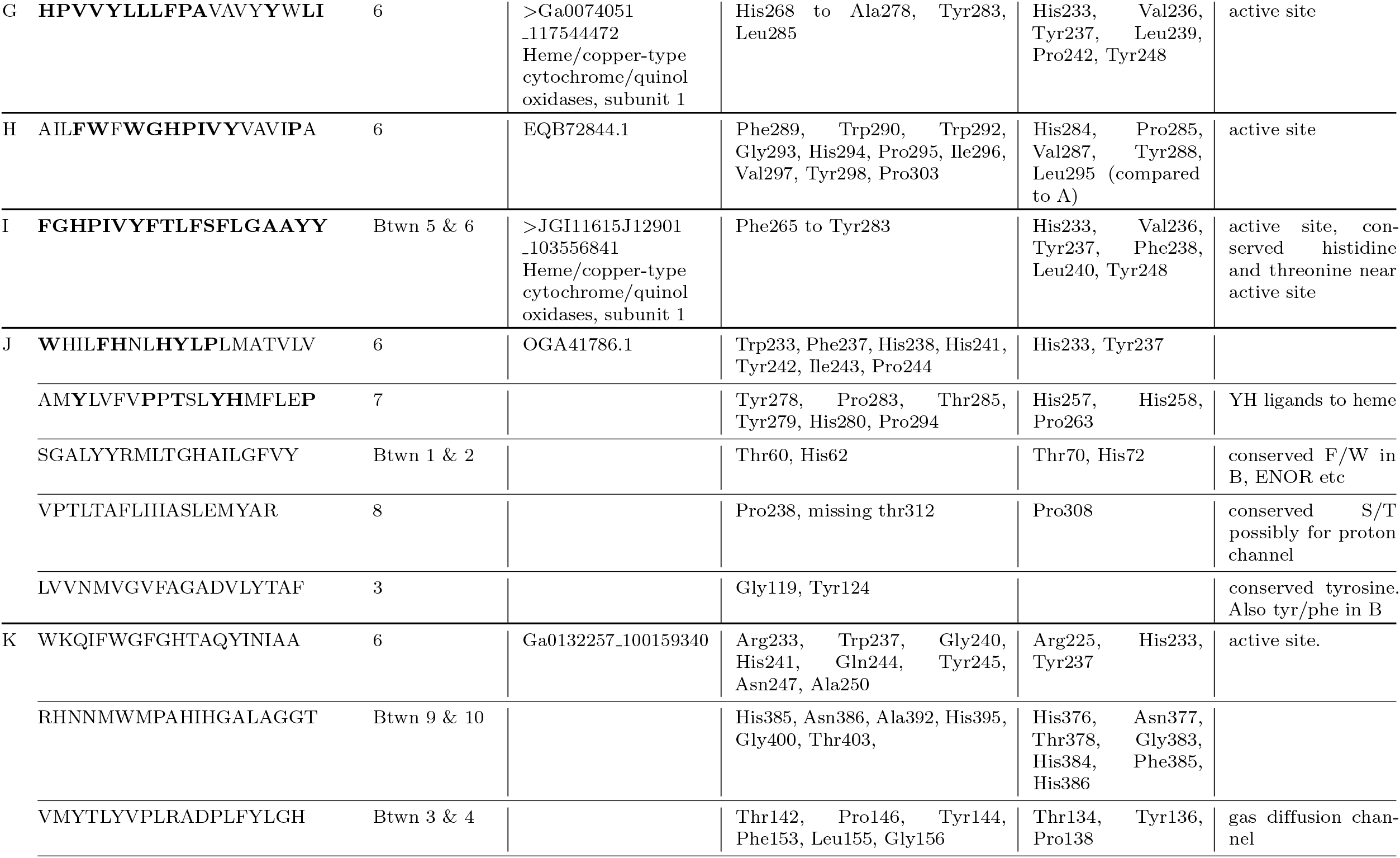
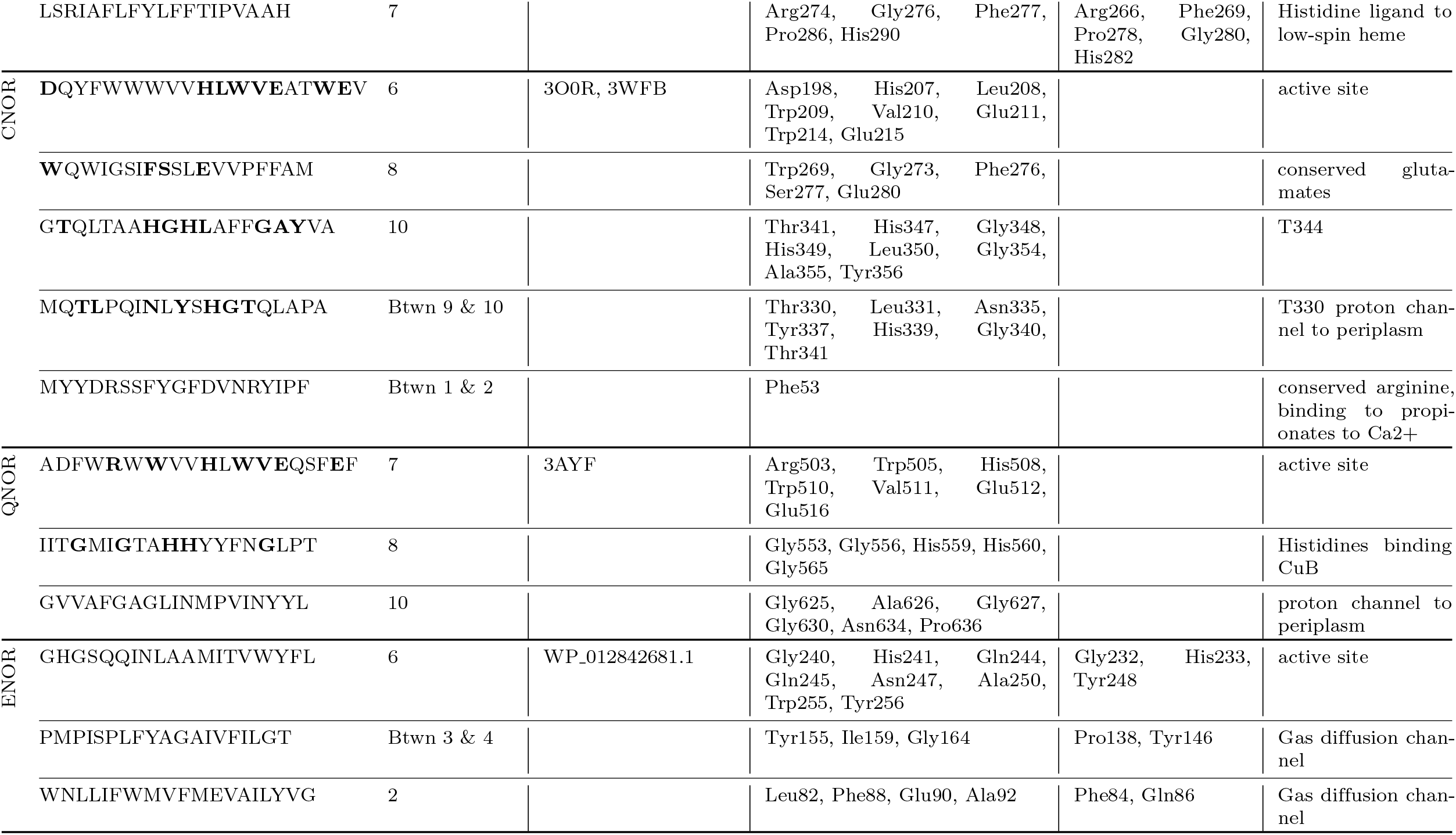
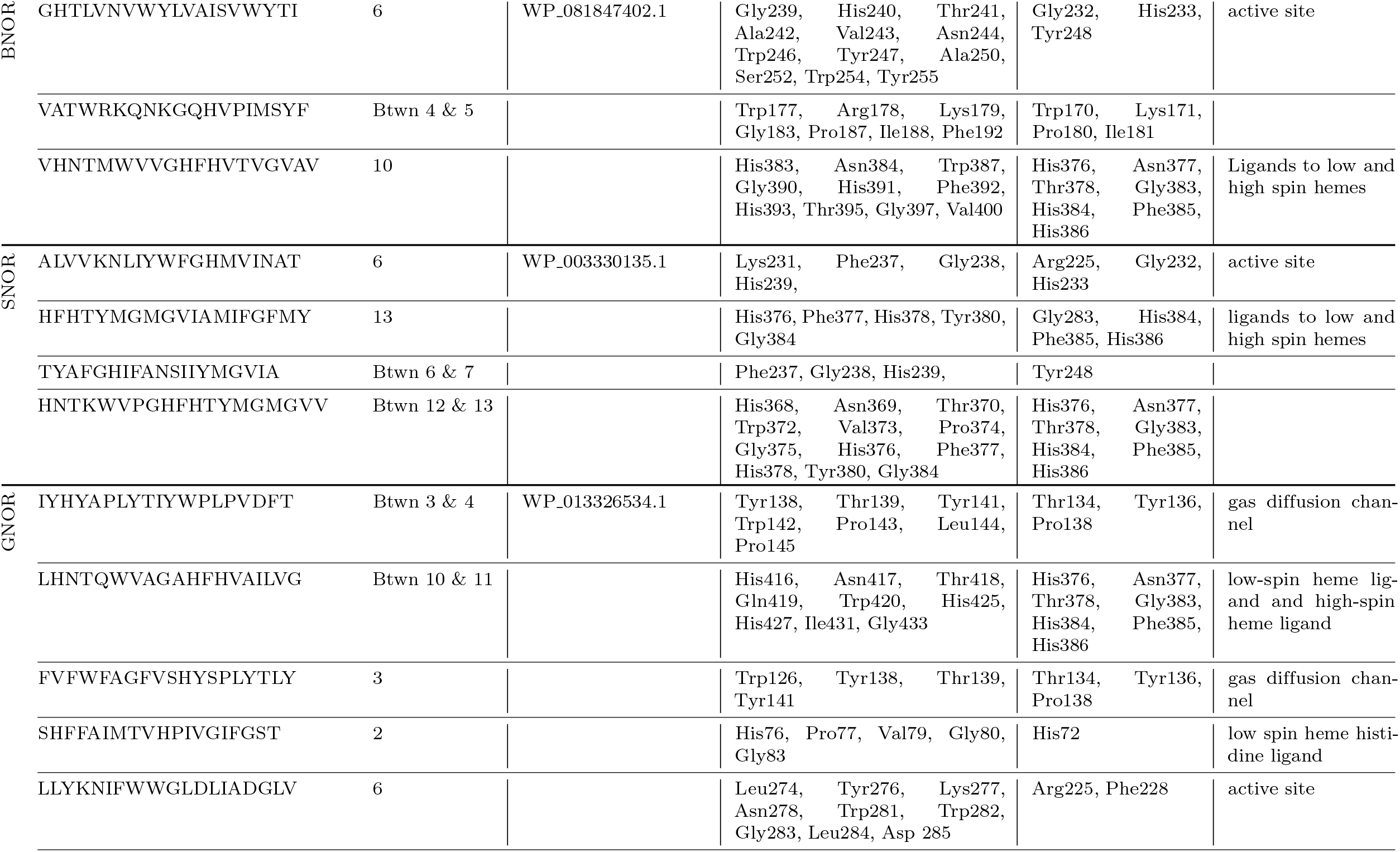
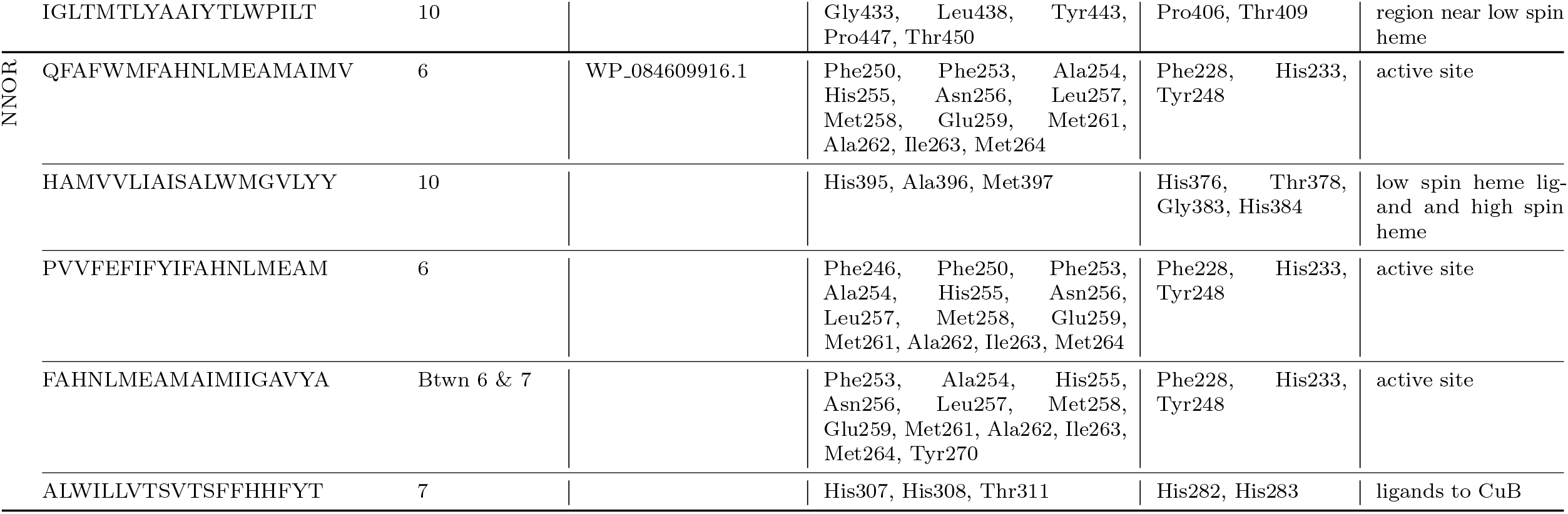
Sequence features discovered by computational model. Amino acid residues conserved within the predictor are indicated as well as their corresponding location in known HCO families.

**Table S2.**
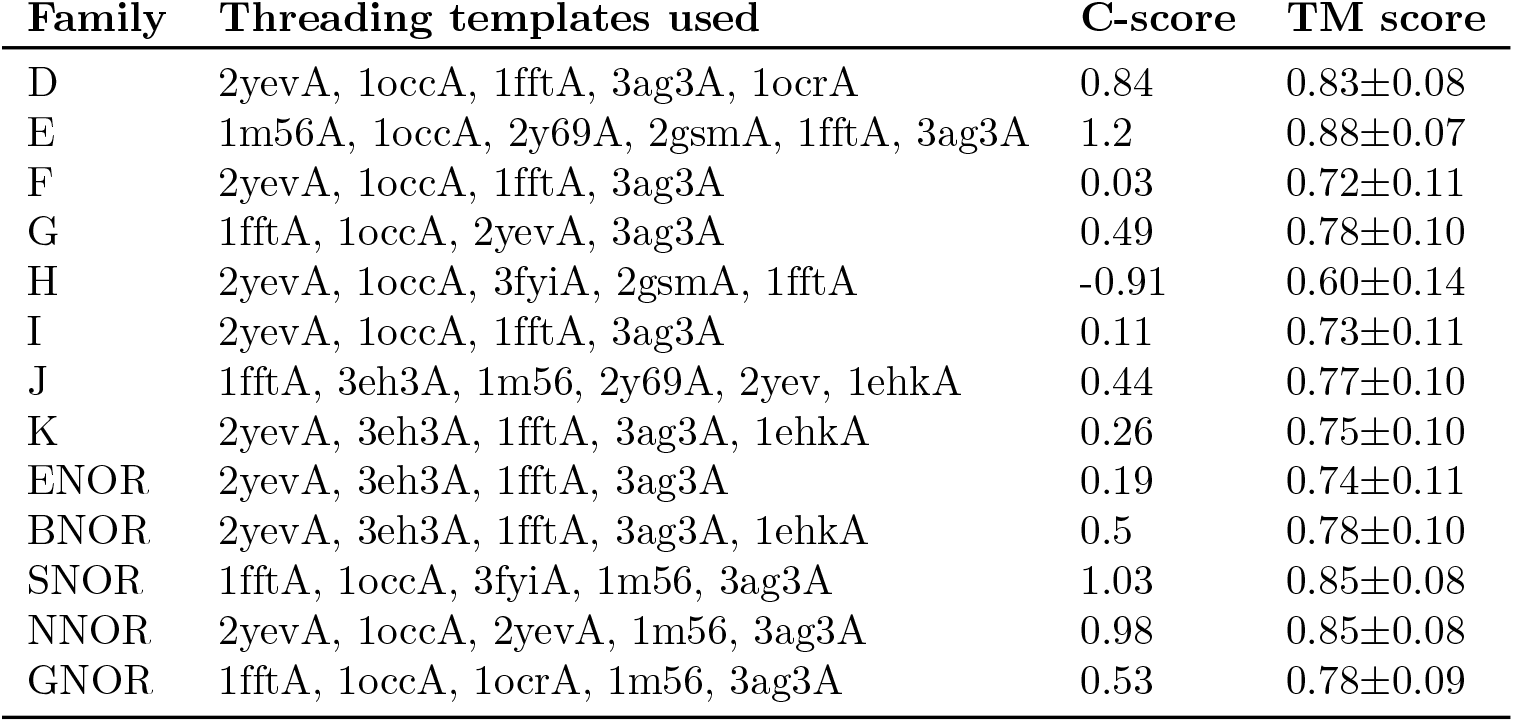
Homology models made for uncharacterized HCOs on i-TASSER with their quality scores

**Fig S1.**
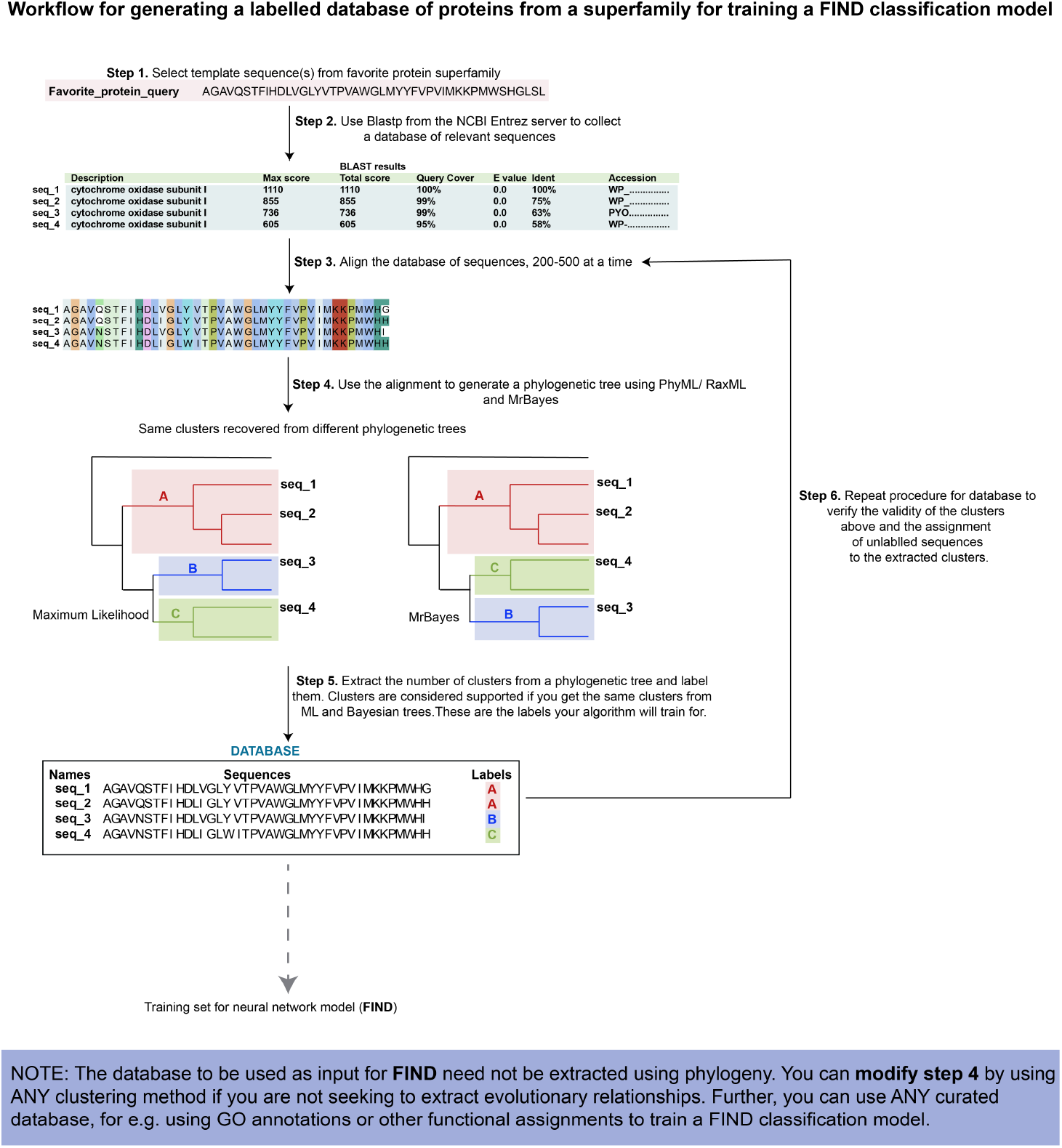
Work flow for the generation of a curated database of protein sequences with labels. This database is then split into training, validation and test set to generate a classification model using **FIND**

**Fig S2.**
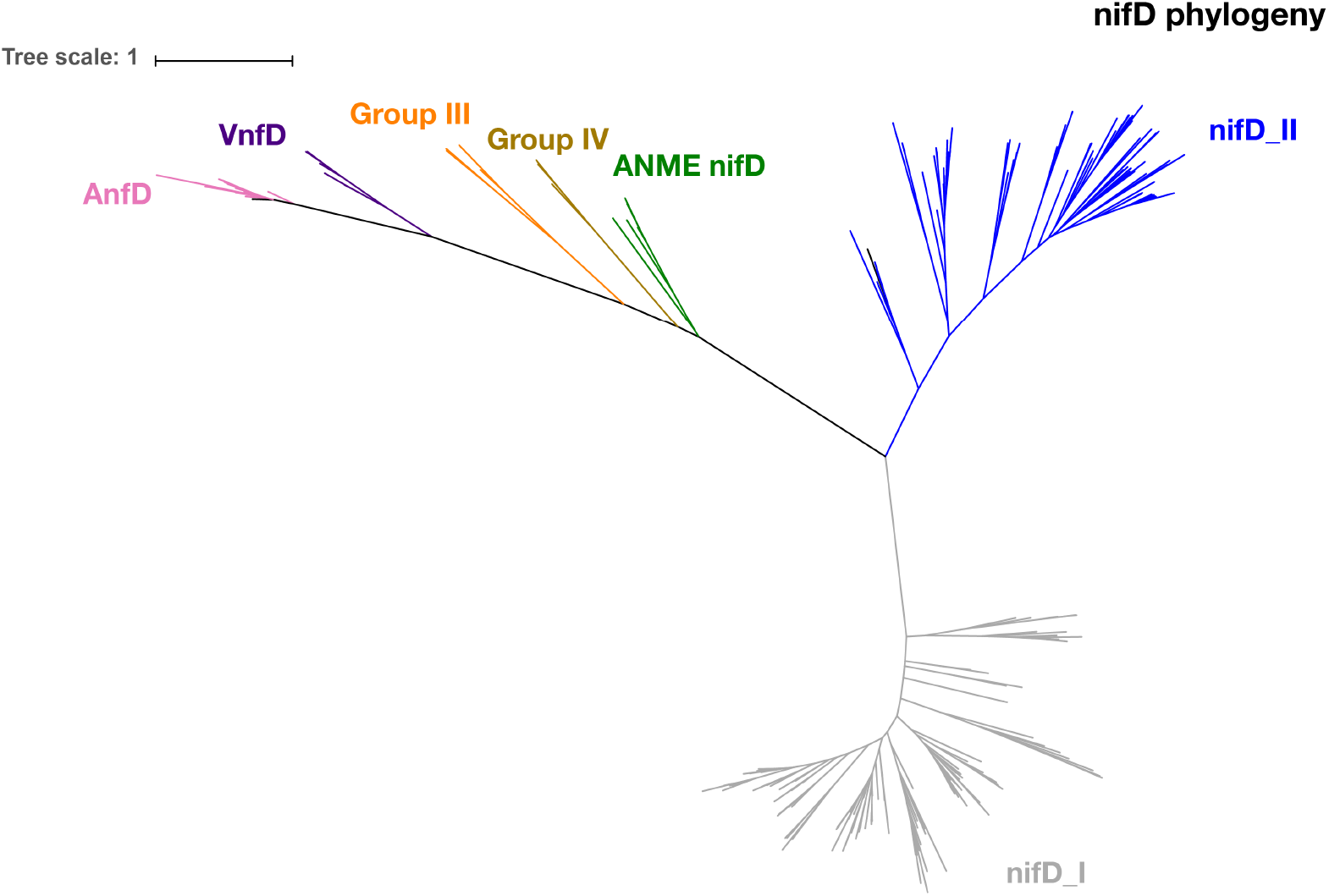
Phylogenetic clustering of nitrogenases using the nifD subunit. Tree was generated using PhyML. Clusters corresponding to nifDI, nifDII, AnfD, VnfD and nifD-ANME were extracted and used to train FIND. Groups III and IV were not used in our training sets because of the smaller number of sequences in those clusters.

**Fig S3.**
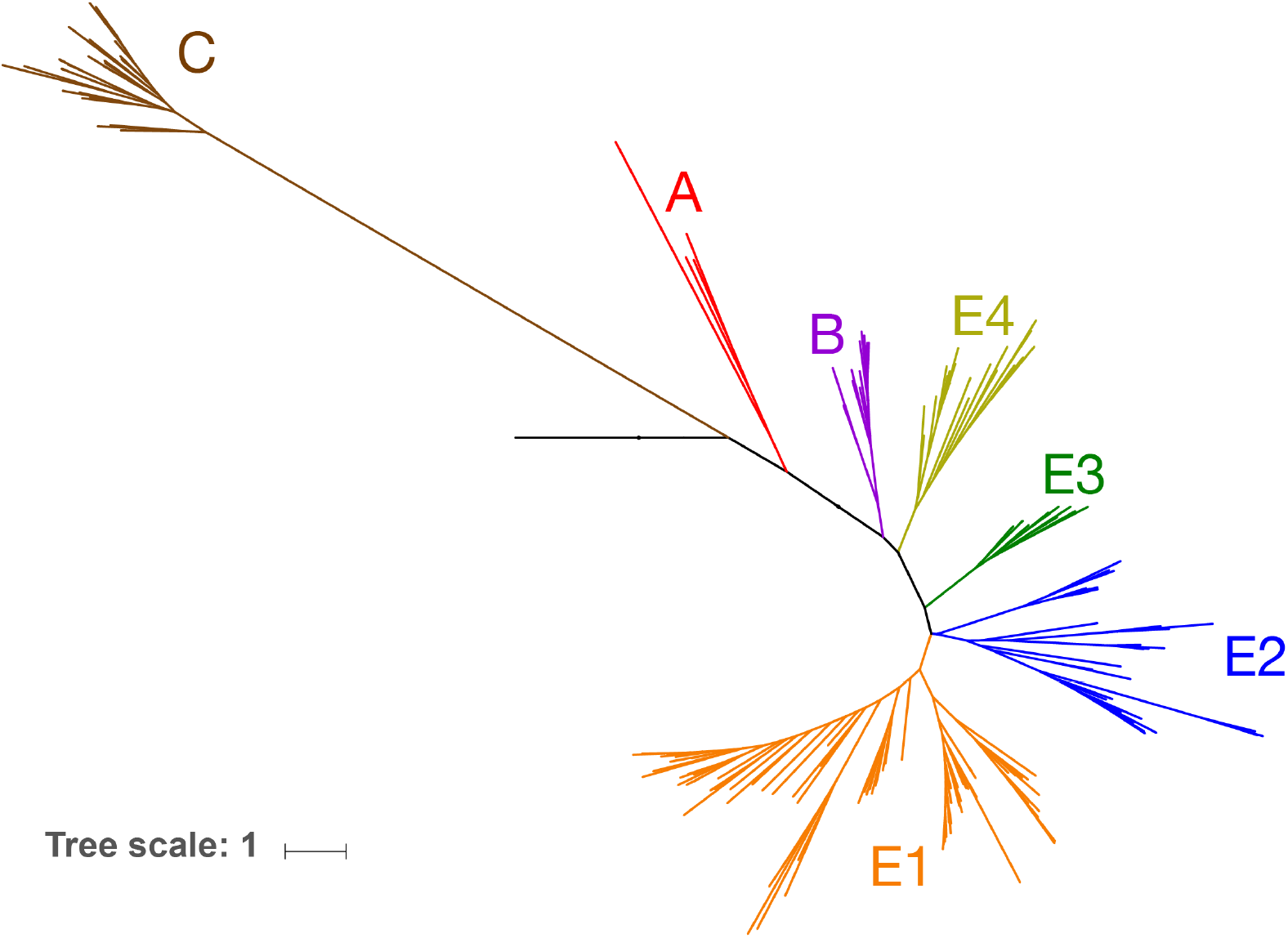
Phylogenetic clustering of cytochrome *bd*-type oxygen reductases using subunit I. This phylogenetic tree was generated using PhyML. Clusters corresponding to E1,E2,E3,E4,B,A and C were extracted and used to train FIND.

## Footnotes

1 previously referred to as the q*Cu_A_*NOR

2 Hemp, Murali publication in preparation

3 http://pytorch.org

4 Hemp, Murali manuscript in preparation

5 communication from Victoria Orphan and Connor Skennerton

6 Pr = tp/(tp + fp), Re = tp/(tp + fn), F1= 2.Pr.Re/(Pr + Re), where tp, tn, fp and fn are true positives, true negatives, false positives and false negatives respectively.

7 Models achieve >90% accuracy with in the first 10 epochs, but may not capture all families until 100. We stopped all runs at a 100 as the learning rate begins to converge to zero

## References

1. Altschul SF, Gish W, Miller W, Myers EW, Lipman DJ. Basic local alignment search tool. Journal of Molecular Biology. 1990;215(3):403–410. doi:https://doi.org/10.1016/S0022-2836(05)80360-2.

2. Clark WT, Radivojac P. Analysis of protein function and its prediction from amino acid sequence. Proteins: Structure, Function, and Bioinformatics. 2011;79(7):2086–2096. doi:10.1002/prot.23029.

3. Altschul SF, Koonin EV. Iterated profile searches with PSI-BLAST—a tool for discovery in protein databases. Trends in Biochemical Sciences. 1998;23(11):444–447. doi:10.1016/S0968-0004(98)01298-5.

4. Sonnhammer ELL, Eddy SR, Durbin R. Pfam: A comprehensive database of protein domain families based on seed alignments. Proteins: Structure, Function, and Bioinformatics. 1997;28(3):405–420. doi:10.1002/(SICI)1097-0134(199707)28:3%A1405::AID-PROT10%BF3.0.CO;2-L.

5. Eddy SR. Profile hidden Markov models. Bioinformatics. 1998;14(9):755–763. doi:10.1093/bioinformatics/14.9.755.

6. Engelhardt BE, Jordan MI, Muratore KE, Brenner SE. Protein Molecular Function Prediction by Bayesian Phylogenomics. PLOS Computational Biology. 2005;1(5). doi:10.1371/journal.pcbi.0010045.

7. Jiang Y, Oron TR, Clark WT, Bankapur AR, D’Andrea D, Lepore R, et al. An expanded evaluation of protein function prediction methods shows an improvement in accuracy. Genome Biology. 2016;17(1):184. doi:10.1186/s13059-016-1037-6.

8. Volpato V, Adelfio A, Pollastri G. Accurate prediction of protein enzymatic class by N-to-1 Neural Networks. BMC Bioinformatics. 2013;14(1):S11. doi:10.1186/1471-2105-14-S1-S11.

9. Szalkai B, Grolmusz V. SECLAF: A Webserver and Deep Neural Network Design Tool for Biological Sequence Classification. ArXiv e-prints. 2017;.

10. Li Y, Wang S, Umarov R, Xie B, Fan M, Li L, et al. DEEPre: sequence-based enzyme EC number prediction by deep learning. Bioinformatics. 2018;34(5):760–769. doi:10.1093/bioinformatics/btx680.

11. Amidi S, Amidi A, Vlachakis D, Paragios N, Zacharaki EI. Automatic single- and multi-label enzymatic function prediction by machine learning. PeerJ. 2017;5:e3095. doi:10.7717/peerj.3095.

12. Song J, Li F, Takemoto K, Haffari G, Akutsu T, Chou KC, et al. PREvaIL, an integrative approach for inferring catalytic residues using sequence, structural, and network features in a machine-learning framework. Journal of Theoretical Biology. 2018;443:125–137. doi:https://doi.org/10.1016/j.jtbi.2018.01.023.

13. Xin F, Myers S, Li YF, Cooper DN, Mooney SD, Radivojac P. Structure-based kernels for the prediction of catalytic residues and their involvement in human inherited disease. BMC Bioinformatics. 2010;11(10):O4. doi:10.1186/1471-2105-11-S10-O4.

14. Petrova NV, Wu CH. Prediction of catalytic residues using Support Vector Machine with selected protein sequence and structural properties. BMC Bioinformatics. 2006;7:312. doi:10.1186/1471-2105-7-312.

15. Hemp J, Gennis RB. In: Schafer G, Penefsky HS, editors. Diversity of the Heme-Copper Superfamily in Archaea: Insights from Genomics and Structural Modeling. vol. 45. Berlin, Heidelberg: Springer Berlin Heidelberg; 2008. p. 1–31.

16. Pereira MM, Sousa FL, Veríssimo AF, Teixeira M. Looking for the minimum common denominator in haem-copper oxygen reductases: Towards a unified catalytic mechanism. Biochimica et Biophysica Acta (BBA)-Bioenergetics. 2008;1777(7):929–934. doi:https://doi.org/10.1016/j.bbabio.2008.05.441.

17. Ettwig KF, Speth DR, Reimann J, Wu ML, Jetten MSM, Keltjens JT. Bacterial oxygen production in the dark. Frontiers in Microbiology. 2012;3:273. doi:10.3389/fmicb.2012.00273.

18. Chang HY, Choi SK, Vakkasoglu AS, Chen Y, Hemp J, Fee JA, et al. Exploring the proton pump and exit pathway for pumped protons in cytochrome ba3 from Thermus thermophilus. Proceedings of the National Academy of Sciences. 2012;109(14):5259–5264. doi:10.1073/pnas.1107345109.

19. Schurig-Briccio LA, Venkatakrishnan P, Hemp J, Bricio C, Berenguer J, Gennis RB. Characterization of the nitric oxide reductase from Thermus thermophilus. Proceedings of the National Academy of Sciences of the United States of America. 2013;110(31):12613–12618. doi:10.1073/pnas.1301731110.

20. Luna VM, Chen Y, Fee JA, Stout CD. Crystallographic Studies of Xe and Kr Binding within the Large Internal Cavity of Cytochrome ba3 from Thermus thermophilus: Structural Analysis and Role of Oxygen Transport Channels in the Heme-Cu Oxidases,. Biochemistry. 2008;47(16):4657–4665. doi:10.1021/bi800045y.

21. Mahinthichaichan P, Gennis RB, Tajkhorshid E. Cytochrome aa3 Oxygen Reductase Utilizes the Tunnel Observed in the Crystal Structures To Deliver O2 for Catalysis. Biochemistry. 2018;57(14):2150–2161. doi:10.1021/acs.biochem.7b01194.

22. Buschmann S, Warkentin E, Xie H, Langer JD, Ermler U, Michel H. The Structure of cbb3 Cytochrome Oxidase Provides Insights into Proton Pumping. Science. 2010;329(5989):327–330. doi:10.1126/science.1187303.

23. Hemp J, Robinson DE, Martinez TJ, Kelleher NL, Gennis RB. The Evolutionary Migration of a Post-Translationally Modified Active-Site Residue in the Proton-Pumping Heme-Copper Oxygen Reductases. Biochemistry. 2006;45(51):15405–15410. doi:10.1021/bi062026u.

24. Hino T, Matsumoto Y, Nagano S, Sugimoto H, Fukumori Y, Murata T, et al. Structural Basis of Biological N2O Generation by Bacterial Nitric Oxide Reductase. Science. 2010;doi:10.1126/science.1195591.

25. Matsumoto Y, Tosha T, Pisliakov AV, Hino T, Sugimoto H, Nagano S, et al. Crystal structure of quinol-dependent nitric oxide reductase from Geobacillus stearothermophilus. Nature Structural &Amp; Molecular Biology. 2012;19:238.

26. Al-Attar S, de Vries S. An electrogenic nitric oxide reductase. FEBS Letters. 2015;589(16):2050–2057. doi:10.1016/j.febslet.2015.06.033.

27. Sievert SM, Scott KM, Klotz MG, Chain PSG, Hauser LJ, Hemp J, et al. Genome of the Epsilonproteobacterial Chemolithoautotroph Sulfurimonas denitrificans. Applied and Environmental Microbiology. 2008;74(4):1145–1156. doi:10.1128/AEM.01844-07.

28. Elman JL. Finding structure in time. Cognitive Science. 1990;14(2):179–211.

29. Hochreiter S, Schmidhuber J. Long Short-Term Memory. Neural Computation. 1997;9(8):1735–1780. doi:10.1162/neco.1997.9.8.1735.

30. Hochreiter S, Bengio Y, Frasconi P, Schmidhuber J. Gradient flow in recurrent nets: the difficulty of learning long-term dependencies. In: Kremer SC, Kolen JF, editors. A Field Guide to Dynamical Recurrent Neural Networks. IEEE Press; 2001. p. 237–243.

31. Bahdanau D, Cho K, Bengio Y. Neural machine translation by jointly learning to align and translate. In: Proceedings of the 3rd International Conference on Learning Representations; 2015.

32. Lecun Y, Boser B, Denker JS, Henderson D, Howard RE, Hubbard W, et al. Backpropagation applied to handwritten zip code recognition. Neural computation. 1989;1(4):541–551.

33. Kingma DP, Ba J. Adam: A Method for Stochastic Optimization. In: Proceedings of the 3rd International Conference for Learning Representations; 2015.

34. Ioffe S, Szegedy C. Batch Normalization: Accelerating Deep Network Training by Reducing Internal Covariate Shift. In: Proceedings of the 32nd International Conference on International Conference on Machine Learning. vol. 37; 2015. p. 448–456.

35. Pettersen EF, Goddard TD, Huang CC, Couch GS, Greenblatt DM, Meng EC, et al. UCSF Chimera—A visualization system for exploratory research and analysis. Journal of Computational Chemistry;25(13):1605–1612. doi:10.1002/jcc.20084.

36. Zhang Y. I-TASSER server for protein 3D structure prediction. BMC Bioinformatics. 2008;9(1):40. doi:10.1186/1471-2105-9-40.

37. Sousa FL, Alves RJ, Pereira-Leal JB, Teixeira M, Pereira MM. A Bioinformatics Classifier and Database for Heme-Copper Oxygen Reductases. PLOS ONE. 2011;6(4):1–9. doi:10.1371/journal.pone.0019117.

38. Adachi S, Nagano S, Ishimori K, Watanabe Y, Morishima I, Egawa T, et al. Roles of proximal ligand in heme proteins: replacement of proximal histidine of human myoglobin with cysteine and tyrosine by site-directed mutagenesis as models for P-450, chloroperoxidase, and catalase. Biochemistry. 1993;32(1):241–252. doi:10.1021/bi00052a031.

39. Zumft WG. Nitric oxide reductases of prokaryotes with emphasis on the respiratory, heme–copper oxidase type. Journal of Inorganic Biochemistry. 2005;99(1):194–215. doi:https://doi.org/10.1016/j.jinorgbio.2004.09.024.

40. Han H, Hemp J, Pace LA, Ouyang H, Ganesan K, Roh JH, et al. Adaptation of aerobic respiration to low O2 environments. Proceedings of the National Academy of Sciences. 2011;108(34):14109–14114. doi:10.1073/pnas.1018958108.

